# The ecological and epidemiological consequences of reproductive interference between the vectors *Aedes aegypti* and *Aedes albopictus*

**DOI:** 10.1101/610451

**Authors:** Robert S Paton, Michael B Bonsall

## Abstract

Vector ecology is integral to understanding the transmission of vector-borne diseases, with processes such as reproduction and competition pivotal in determining vector presence and abundance. The arbovirus vectors *Aedes aegypti* and *Aedes albopictus* compete as larvae, but this mechanism is insufficient to explain patterns of coexistence and exclusion. Inviable interspecies matings - known as reproductive interference - is another candidate mechanism. Here, we analyse mathematical models of mosquito population dynamics and epidemiology which include two *Aedes*-specific features of reproductive interference. First, as these mosquitoes use hosts to find mates, reproductive interference will only occur if the same host is visited. Host choice will, in turn, be determined by functional responses to host availability. Second, females can become sterilised after mis-mating with heterospecifics. We find that a species with an affinity for a shared host will suffer more from reproductive interference than a less selective competitor. Costs from reproductive interference can be “traded-off” against costs from larval competition, leading to competitive outcomes difficult to predict from empirical evidence. Sterilisations of a self-limiting species can counter-intuitively lead to *higher* densities than a competitor suffering less sterilisation. We identify that functional responses and reproductive interference mediate a concomitant relationship between vector ecological dynamics and epidemiology. Competitors with opposite functional responses can maintain disease where human hosts are rare, due to vector coexistence facilitated by a reduced cost from reproductive interference. Our work elucidates the relative roles of the competitive mechanisms governing *Aedes* populations and the associated epidemiological consequences.

## 1. Background

An estimated 390 million annual cases of dengue and the emergence of Zika has motivated policy makers, NGOs and academics to call for the management of vector populations (Bhatt et al., 2013; Yakob and Walker, 2016). Effective disease mitigation requires that vector occurrence and abundance can be accurately predicted (Bhatt et al., 2013), making it essential that we understand the ecological processes driving the distributions of the principal vectors of Zika, dengue and chikungunya - *Aedes aegypti* and *Aedes albopictus*.

*Ae. aegypti* is widely considered the primary vector of flaviviruses (Black et al., 2002; Chouin-Carneiro et al., 2016), with a propensity for biting and living in proximity to humans (Scott and Takken, 2012). Native to Africa, *Ae. aegypti* has become established across Asia and the Americas (Kraemer et al., 2015). In contrast, *Ae. albopictus* is considered by some to be a secondary vector, but is still capable of transmitting over 26 different pathogens (Paupy et al., 2009). It originated from South-East Asia (Gratz, 2004) and is better able to colonise temperate environments relative to *Ae. aegypti* (Kraemer et al., 2015). *Ae. albopictus* is considered an opportunistic feeder (Paupy et al., 2009; Richards et al., 2006), however in some locations it has been shown it to be anthropophilic (Ponlawat and Harrington (2005); Delatte et al. (2010)).

Human activities, such as international shipping, have expanded the distributions of *Ae. aegypti* and *Ae. albopictus* (Lounibos and Juliano, 2018), leading to the species coming into contact as they establish in new regions. They share a similar ecological niche, with females laying eggs in small (often ephemeral) pools of water which hatch into a mobile larval stage. Early introductions of *Ae. aegypti* to Asia resulted in the displacement of *Ae. albopictus* in places such as Bangkok and Kuala Lumpur, but more recently *Ae. albopictus* rapidly displaced *Ae. aegypi* from all but urban areas in the Southern United States (O’Meara et al., 1995; Lounibos and Juliano, 2018). Both Braks et al. (2004) and Simard et al. (2005) document co-occurrence in Brazil and Cameroon respectively, albeit in segregated habitats. Habitat segregation is supported by (Rey et al., 2006), who found *Ae. aegypti* to have a preference for urban landscapes and *Ae. albopictus* for suburban areas with more vegetation. It is widely accepted that these species compete in the wild, leading to different patterns of coexistence and competitive exclusion (Lounibos and Juliano, 2018).

The most obvious stage for this competition is as larvae, where individuals will compete for food and space. The importance of density-dependent larval-stage competition in regulating *Aedes* populations has long been recognised (Southwood et al., 1972; Dye, 1984), but the exact strength and form is difficult to quantify in wild populations (Legros et al., 2009). This has led researchers to investigate populations of mosquitoes in lab or semi-field settings (many are reviewed in Juliano (2009)). Most studies rear populations of *Ae. aegypti* and *Ae. albopictus* in different density conditions and assess the penalties in survival and development time. In 2010, Juliano conducted a meta-analysis of larval competition studies, finding that the magnitude and sign of inter-specific larval competition depends on the resources that the larvae were reared on. Subsequent studies (e.g. Reiskind et al. (2012)) have supported this resource-dependent variation in competitive outcome. Leisnham et al. (2009) found that this context-dependence can also be observed between spatially-distinct mosquito populations. In the absence of a consistent rule for the competitive exclusion and coexistence of these disease vectors, another mechanism has increasingly been explored: reproductive interference.

Reproductive interference occurs when heterospecifics engage in mating activities (e.g. courtship, copulation) that reduce the fitness of one or both parties (Ribeiro and Spielman, 1986; Burdfield-Steel and Shuker, 2011). It has been documented in a range of different species, including arthropods (Kishi et al., 2009; Shuker et al., 2015). In their review, Gröning and Hochkirch (2008) describe seven types of reproductive interference; signal jamming, heterospecific rivalry, misdirected courtship, heterospecific mating attempts, erroneous female choice, heterospecific mating, and hybridisation.

Erroneous mate selection is known to occur in *Aedes*. Tripet et al. (2011) found that non-viable interspecies matings between *Ae. aegypti* and *Ae. albopictus* occurred at low rates in the field when screening for heterospecific sperm. In lab experiments, Bargielowski et al. (2013) found far higher rates than were observed in the field, as well as a propensity for *Ae. aegypti* females to be more likely to mis-mate than *Ae. albopictus*. In the same experiment, *Ae. aegypti* strains taken from populations which have not encountered *Ae. albopictus* (allopatric) mis-mated more frequently than those from populations sympatric with the competitor, suggesting an asymmetric penalty in reproductive interference that may be selecting for more discerning *Ae. aegypti* females in the wild. *Ae. aegypti* was similarly found to mis-mate more often in lab experiments by Marcela et al. (2015). While the higher rates found in the lab will be at least partially attributable to “cage effects” - whereby females are harassed due to crowding - it is possible that there is a true discrepancy in interspecies mating rates in the field. These interspecies matings do not produce viable offspring, and incur a penalty in wasted courtship/handling time and fitness costs to females through harassment. From this evidence, it is clear that at least four of the seven forms of reproductive interference that Gröning and Hochkirch (2008) describe are found in *Aedes*: misdirected courtship, heterospecific mating attempts, erroneous female choice and heterospecific mating. All of these factors will contribute to a diminished reproductive rate when in the presence of heterospecifics. We do not include signal interference here, but it is well understood that the harmonisation of harmonic wing-beat frequencies of *Ae. aegypti* males and females increases copulation success (Cator et al., 2009), a process which is yet to be examined between *different Aedes* species.

*Ae. albopictus* females mis-mated to *Ae. aegypti* males will re-mate with a higher frequency than in the reverse case, with *Ae. aegypti* females sometimes becoming refractory to further mating attempts (Tripet et al., 2011). We refer to this effect as sterilisation. Moreover, sterilisation can be induced in *Ae. aegypti* females solely by the injection of *Ae. albopictus* accessory gland proteins, without the successful transfer of sperm (Carrasquilla et al., 2015). Sterilisation could therefore be induced more widely than is typically observed when screening only for the presence of heterospecific sperm in spermatheca. Should female mosquitoes become refractory in the wild, these mis-mated females will not contribute to the larval population, consequently diminishing the regulatory impact of density-dependence.

Unlike *Anopheline* mosquitoes which mate in crepuscular swarms, *Aedes* mosquitoes use hosts as beacons to swarm and find mates (Nelson, 1986; Yuval, 2006). However, not all vectors respond in the same way to host availability, with some showing generality and others a high degree of specificity. This means that the overlap in the functional responses of *Ae. aegypti* and *Ae. albopictus* will scale the number of heterospecifics that will be encountered, as the process of reproduction is mediated by host choice. It is has been documented that *Ae. albopictus* can have plastic responses to host availability (Valerio et al., 2010), with *Ae. aegypti* more consistently anthropophilic (Richards et al., 2006; Paupy et al., 2009).

Any model of *Aedes* reproduction would therefore need to include a flexible function to describe how the encounter rate between the two species changes with the availability of shared hosts. The epidemiological importance of functional responses to host availability was explored in detail by Yakob (2016), where a function capable of describing the range of host selection processes exhibited by biting insects was developed. Using a mathematical model, Yakob (2016) was able to show that host functional responses can have profound effects on the epidemiology of vector-borne pathogens, determining the level of control required to prevent transmission. We link this same host functional response to vector population dynamics; host choice will both determine patterns of vector coexistence and exclusion, while also altering biting rates and changing transmission.

Asymmetric reproductive interference is now considered by many to be an important driver of coexistence and competitive exclusion in *Aedes*, particularly in the absence of a consistent rule for the outcome of larval competition (Bargielowski and Lounibos, 2016; Lounibos and Juliano, 2018). Studies on *Aedes* have hitherto had to rely on *general* models of reproductive interference to make inference on the relative roles of reproductive interference and larval competition (Lounibos and Juliano, 2018). For instance, Kuno (1992) presented a theoretical model of the combined effects of mating interference and standard Lotka-Volterra competition (later revisited in detail by Kishi and Nakazawa (2013)). The model includes a frequency-dependent function which scales the reproductive rate of a species with increasing numbers of heterospecifics. They demonstrate that the process of reproductive interference can diminish the possibility of coexistence and introduce priority effects, which make invasions challenging. This model has provided great insight into this process, but does not include the *Aedes*-specific features described above; namely that encounter rates will vary with host availability, and females could become refractory after heterospecifc matings. In this study, we use a mathematical model to explore these two aspects of *Aedes* reproductive biology. First, we examine how functional responses to host availability could alter the encounter rate of heterospecific mosquitoes and, in turn, the feasibility of coexistence. Second, we consider how different ratios of females becoming sterilised after heterospecific matings can lead to different patterns of coexistence and exclusion. We conclude by examining the consequences of these ecological processes for vector-borne diseases.

## 2. Methods

In this research we build two mathematical models of *Aedes* population dynamics; first we include functional responses to host availability, then female sterilisation by heterospecific copulations. We do not assign either *Ae. aegypti* or *Ae. albopictus* to a particular state variable and instead discuss our findings in the context of species “*A*” and “*B*”. This allows us to draw overarching conclusions about the nature of the processes we examine without using species-specific model parameterisations from particular lab or field studies, which would risk obfuscating conclusions. Shared parameters (such as birth and death rates) are therefore taken to be of a “general” *Aedes* mosquito.

We proceed to add a model of vector-borne disease dynamics to our population dynamics model. This epidemiological model used parameters from the literature on dengue. Again, we use the same parameters across both species so that conclusions can be drawn about the processes of interest.

### 2.1 Ecological Models

Kuno (1992) presented a model of reproductive interference which was further analysed by Kishi and Nakazawa (2013):

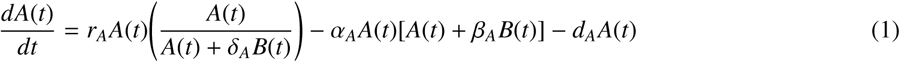

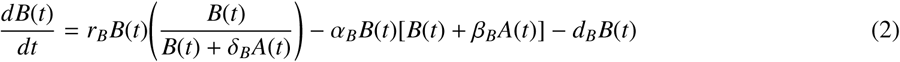

where *A* and *B* are the densities of the *i*^*th*^ species, with *r*_*i*_ the reproductive rate, *d*_*i*_ the death rate, α_*i*_ the strength of intraspecific competition, β_*i*_ the *relative* strength of interspecific competition and α_*i*_ the strength of reproductive interference. The parameters for reproductive interference, δ_*i*_, is part of a frequency-dependent function describing the proportional reduction in reproductive rate with increasing densities of heterospecifics. Equations 1 and 2 have two single-species dominant roots which are stable 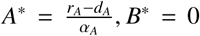 and 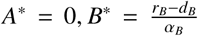, and an unstable α_*A*_α_*B*_trivial (*A*^*^ = 0, *B*^*^ = 0) root. Should interspecific competition exceed intraspecific competition, there will be one unstable multi-species root. If intraspecific is greater than interspecific, then there will be one stable coexistence point and two unstable coexistence points (Kuno, 1992; Kishi and Nakazawa, 2013). These unstable “saddle-points” are caused by regions of the phase-space where the effects of interspecific competition outweigh intraspecific competition, leading to an *Allee* effect. This manifests as a priority effect, whereby the frequency-dependent costs of reproductive interference make it difficult for a species to invade from rare as it suffers a large penalty in reproductive interference from a more abundant competitor.

#### 2.1.1 Functional responses to hosts

In order to include host functional responses, the model had to reflect that heterospecifics will only be encountered by mosquitoes utilising the same hosts:

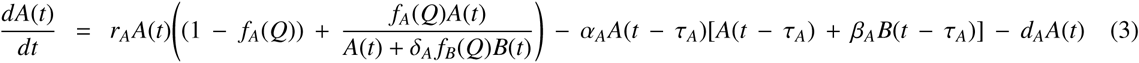

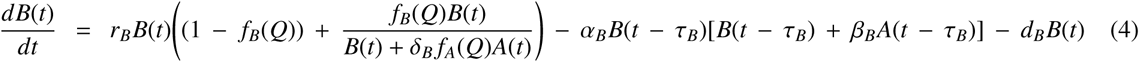

This form is similar to Kuno’s (1992) model but has three key modifications. First, the model is now time-lagged to represent the delay in the adult recruitment due to the larval stage of *Aedes* mosquitoes. Therefore larvae of the *i*^*th*^ species experience density-dependent intra-and interspecific competition at time *t* − τ_*i*_, while female egg laying and adult deaths occur in the present time. Second, the parameter β_*i*_ now corresponds to the relative strength of interspecific *larval* competition in relation to intraspecific competition (α_*i*_). Third, the process of reproductive interference is now mediated by the functional response to host availability (taken from Yakob (2016), inspired by Real (1977)), through the function *f*_*i*_(*Q*):

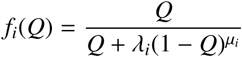

where *Q* (*Q* ∈ [0, 1]) is the proportion of the shared host in relation to all other hosts, and *λ*_*i*_ and *µ*_*i*_ are the shape parameters which describe how the *i*^*th*^ species utilises the shared host. A description of these parameters and an illustration of the scenarios explored here are given in Table 1. These functions ensure that only the fraction of the population using a shared host will be exposed to heterospecifics and suffer from reproductive interference. This allows the proportion of successful matings to be described as:

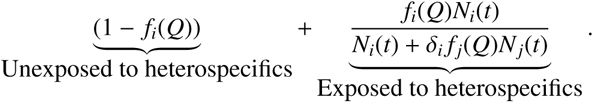

**Table 1:**
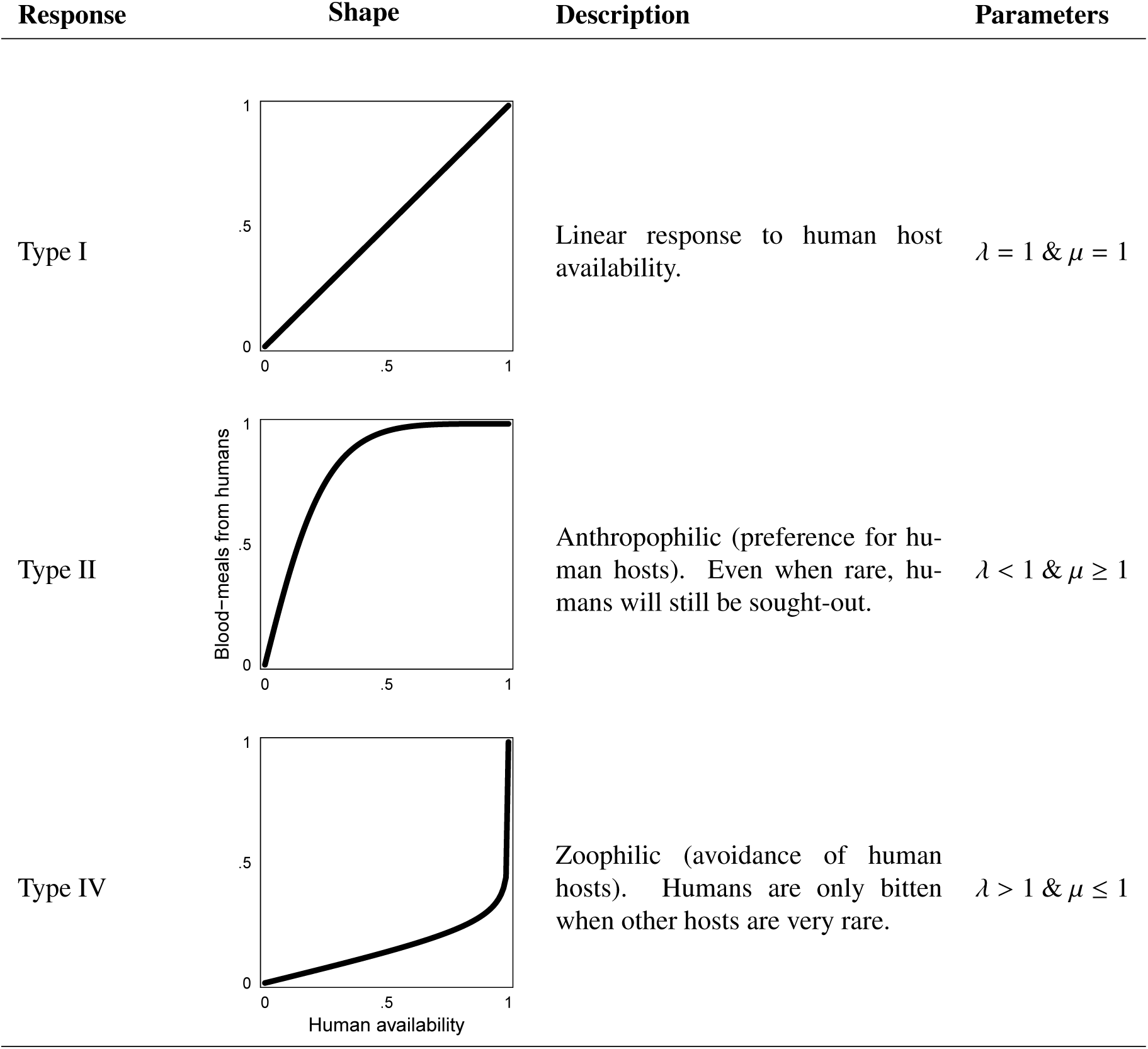
Table of the of functional responses explored in this paper using Yakob’s (2016) function, *f* (*Q*) = *Q*/(*Q* + λ(1 *Q*)^µ^). The proportional availability of human hosts, *Q*, is shown on the x-axis while the proportion of blood-meals taken from human hosts (*f* (*Q*)) is on the y-axis. Other functional forms are possible, but for conciseness we only draw the distinction between linear, anthropophilic and zoophilic.

| Response | Shape | Description | Parameters |
| --- | --- | --- | --- |
| Type I | | Linear response to human host availability. | $\lambda = 1 \text{ \& } \mu = 1$ |
| Type II | | Anthropophilic (preference for human hosts). Even when rare, humans will still be sought-out. | $\lambda < 1 \text{ \& } \mu \geq 1$ |
| Type IV | | Zoophilic (avoidance of human hosts). Humans are only bitten when other hosts are very rare. | $\lambda > 1 \text{ \& } \mu \leq 1$ |

The first part of this expression gives the proportion of the population of the *i*^*th*^ species which is not utilising the shared host and will therefore not suffer from reproductive interference. Note that, like epidemiological models of vector-borne pathogens, we assume that inter-or intraspecific competition will *not* impede access to the host, as the host is many orders of magnitude larger than the vector. The unexposed term will tend to zero when the shared host is the only host available (*Q* → 1 therefore *f*_*i*_(*Q*) → 1). In the second part, the numerator is the number of the *i*^*th*^ species using the shared host, and the denominator the sum of conspecifics and heterospecific matings with individuals of the *j*^*th*^ species. In our model, we make the simplifying assumption that the shared host is a human, and that the mosquitoes will not encounter each other at other vertebrate hosts. As described in Table 1, we limit our functional responses to linear, anthropophilic and zoophilic. While both species are known to bite other animal hosts, they do so with different preferences which vary between species and strain. Therefore, the “true” functional response would be a complex aggregate of the responses to *all* shared hosts, and maybe different for different mosquito *strains*.

#### 2.1.2 Sterilised Females

Equations 1 and 2 were also modified to reflect that some female *Aedes* do not go on to find a conspecific mate after mating with a heterospecifics. Sterilisation can be modelled by scaling the strength of density-dependent larval competition by the fraction of females contributing eggs (and therefore larvae) to shared pools of water:

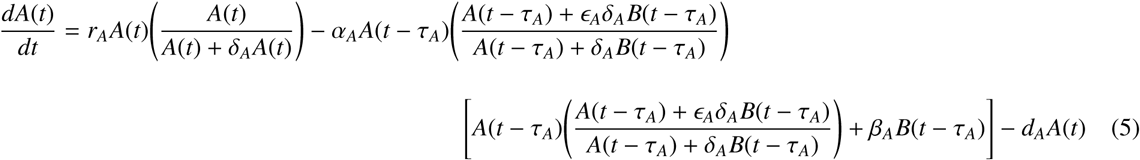

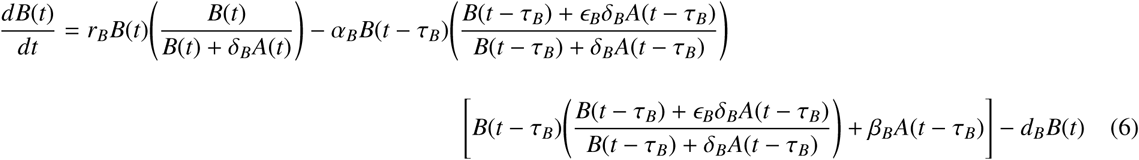

The terms describing the larval stage density-dependent interactions are now scaled by the effects of reproductive interference (as refractory females will not lay eggs in the future). The parameter E_*i*_ is the *proportion* of females of the *i*^*th*^ species that *do not* become refractory after a copulation with a heterospecific male, and contribute larvae to the next generation (E_*i*_ ∈ [0, 1]).

### 2.2 Epidemiological Models

We were interested in examining the implications of our ecological models on the epidemiology of a theoretical vector-borne pathogen. It was necessary for this model to include how functional responses to host availability alters biting rates (akin to the system explored by Yakob (2016)). The transmission of a pathogen in this two-vector system can therefore be described by modifying the Ross-MacDonald model of vector-borne disease transmission:

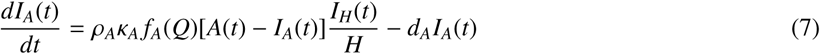

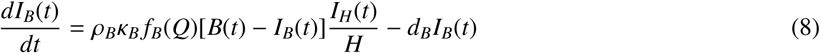

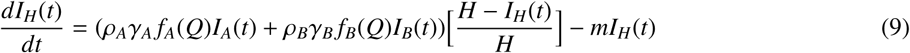

The state variables *I*_*A*_ and *I*_*B*_ are the density of infected mosquitoes of species *A* and *B*, while *I*_*H*_ is the density of infected hosts. The parameter ρ_*i*_ is the biting rate (scaled by the functional response to hosts *f*_*i*_(*Q*)), κ_*i*_ the rate at which bites on infected hosts lead to an infection and γ_*i*_ the rate at which bites from an infected mosquito infecting a human for the *i*^*th*^ mosquito species. *H* is the *fixed* density of hosts in the system and *m* the rate of recovery of hosts. The death rate of infected mosquitoes is given by *d* (such that infected mosquitoes die at an equal rate to uninfected mosquitoes and do not recover from infection). Parameter descriptions and values are given in Table 2.

**Table 2:** Table of parameter values examined in this paper. Sub-scripted parameters indicate that these were varied asymmetrically between the two species. Otherwise, they were always the same for both.

| Parameter | Definition | Value | Reference |
| --- | --- | --- | --- |
| **Ecological parameters** |  |  |  |
| $r$ | Reproductive rate | 1.31 | Southwood et al. (1972); Dye (1984) |
| $d$ | Death rate | 0.12 | Southwood et al. (1972); Dye (1984) |
| $\alpha$ | Strength of density-dependent (intraspecific) larval competition | 1 | - |
| $\beta_i$ | Relative strength (in relation to intraspecific) of interspecific larval competition | varied | - |
| $\delta_i$ | Strength of reproductive interference | varied from 0 to 1 | Kishi and Nakazawa (2013) |
| $\lambda_i, \mu_i$ | Shape parameters of the functional response | see table 1 | Yakob (2016) |
| $Q$ | The proportional availability of human hosts | varied | Yakob (2016) |
| $\epsilon_i$ | The proportion of females that contribute larvae to the next generation after interspecific matings | varied from 0 to 1 | - |
| $\tau$ | Time from an egg to adult emergence | Assumed 0 in simulations | - |
| **Epidemiological parameters** |  |  |  |
| $\rho$ | The biting rate of adult mosquitoes | 0.5 | Alphey et al. (2011) |
| $\kappa$ | The probability of a bite from an infected mosquito converting into an infection in a human | 0.38 | Alphey et al. (2011) |
| $\gamma$ | The probability of a bite on an infected human converting into an infection in the mosquito | 0.38 | Alphey et al. (2011) |
| $m$ | The recovery rate of infected humans | 0.17 | Alphey et al. (2011) |
| $H$ | The density of human hosts | varied | - |

### 2.3 Model analysis

All mathematical derivations were confirmed in the software Mathematica (version 11.3.0.0) (Wolfram Research, 2018) and all calculations and simulations were conducted in the statistical software R (version 3.5.1) (R Core Team, 2016). Code has been made available as supplementary material online.

#### 2.3.1 Boundaries for coexistence and isoclines

We solved equations 3 and 4 for *dA*(*t*)/*dt* = 0 and *dB*(*t*)/*dt* = 0 to give two quadratic expressions (see the appendix). The positive solution of these expressions described the zero-net-growth isoclines for the system. Substituting the solution for species *A* into the solution for species *B*, and vice-versa, yielded two cubic expressions for which the three roots are the potential equilibrium population sizes of the respective species. The discriminant of these expressions reveals the nature of these roots; if the discriminant of both is greater than 0, then there are three non-zero real roots (one stable flanked by two unstable) and coexistence is possible (Kishi and Nakazawa, 2013). Otherwise there is an *unstable* coexistence point, where coexistence is unsustainable. By examining the regions of the parameter space where these conditions are met, we can examine the requisite conditions for coexistence. These expressions are given in the appendix and the supplementary Mathematica file. We also derived cubic isoclines for equations 5 and 6. For both sets of isoclines, the stability of the equilibria was determined from the dominant eigenvalue of the Jacobian matrix of the linearised system, which is given in the supplementary Mathematica file.

#### 2.3.2 Deriving the basic reproductive number, R_0_

The equations 7, 8 and 9 can be used to determine the basic reproductive number of the disease, *R*_0_, in a system where vector populations are at equilibrium. If both mosquito populations are at equilibrium (*A*(*t*) = *A*^*^, *B*(*t*) = *B*^*^), and the infected populations tend to zero (*I*_*A*_(*t*), *I*_*B*_(*t*) and *I*_*H*_(*t*) → 0) we can derive a Jacobian matrix using equations 7 to 9 to determine the stability of this disease-free state:

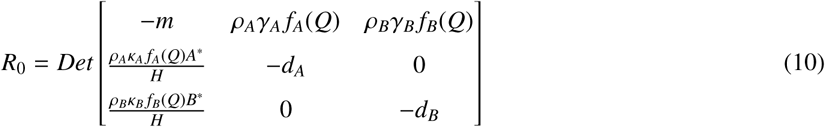

When the determinant of this matrix is less than zero, then the disease-free equilibrium is stable. If the determinant is greater than zero then it is unstable. A full derivation is given in the appendix.

#### 2.3.3 Stochastic simulations

Stochastic simulations demonstrate how a system of differential equations respond to random perturbations. In our case, this is particularly interesting given that our system yields multi-stable population dynamics; determining the asymptotic stability *precisely* at an equilibrium does not fully communicate what the *global* stability of this equilibrium is relative to other states (Nolting and Abbott, 2016). For instance, transitions between asymptotically stable equilibrium can occur when stochastic perturbations “knock” the state of the system beyond the domain of attraction of one equilibrium into the domain of another. This can demonstrate the *relative* stability of equilibrium in multi-stable systems.

The delay-differential equations given above (3 to 9) are of the form 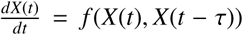 We can perturb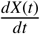using discrete increments (*dW*) of a Wiener random walk process (*W*) where each *dW* is taken from a normal distribution with a zero mean and variance *dt* (Higham, 2001). For our multi-variate example (species *A* and *B* and infecteds), the differential equations can be re-written in the form:

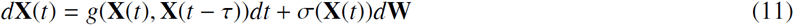

where **X**(t) is a column vector of *N* state variables at time *t*, **X**(*t* - τ) are the time-lagged state variables, and **W** a vector of *N* independent Weiner processes (one for each variable). The function *g* is the deterministic component of the processes described by equations 3 through 9, while the function σ describes the way in which the Weiner process acts on the state variables (Higham, 2001). Note that the term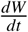 is absent here as the Wiener process is almost always not differentiable. The function σ(**X**(*t*)) describes how the increments of noise (*d***W**) impact each of the state variables. We assume that the function σ is the same for each state variable;

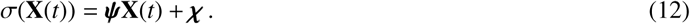

The effect of demographic noise (as a function of the the current population density) for each state-variable is described by the parameter vector **ψ**, while exogenous sources of noise (e.g. migration) are described by the vector of parameters **χ**. Numerical simulations were conducted using a Euler-Maruyama scheme, as outlined in Higham (2001).

## 3. Results

### 3.1 Ecology

The strength of reproductive interference changes the region of parameter space in which coexistence is possible. In Figure 1, this is shown across different combinations of functional responses (linear, anthropophilic or zoophilic) in the rows and host availability in the columns. In all cases, increasing the cost of reproductive interference (δ_*i*_) reduces the possibility of coexistence. Reducing host availability (*Q*) expands the region where coexistence is possible. This expansion is symmetric when both species share a linear functional responses in panels A to C. However, when the species have different functional responses and humans are not the only available host (*Q* < 1), the region of coexistence changes asymmetrically (observed in panels E, F, H, I, K and L). In these asymmetric cases, the species which is disinclined to use a shared host can suffer a greater cost of reproductive interference while still coexisting with an anthropophilic competitor. This is most pronounced in the panels E, F, K and L. Panels H and I give are the only cases where the region of coexistence is smaller in comparison to panels B and C. This occurs as encounter rates in this scenario are *higher* than the linear/linear default case. Increases/decreases in *β*_*i*_ simply shrink/grow the overall area of the coexistence region of while retaining the same overall patterns (not shown, analogous to Kishi and Nakazawa (2013)).

**Figure 1:**
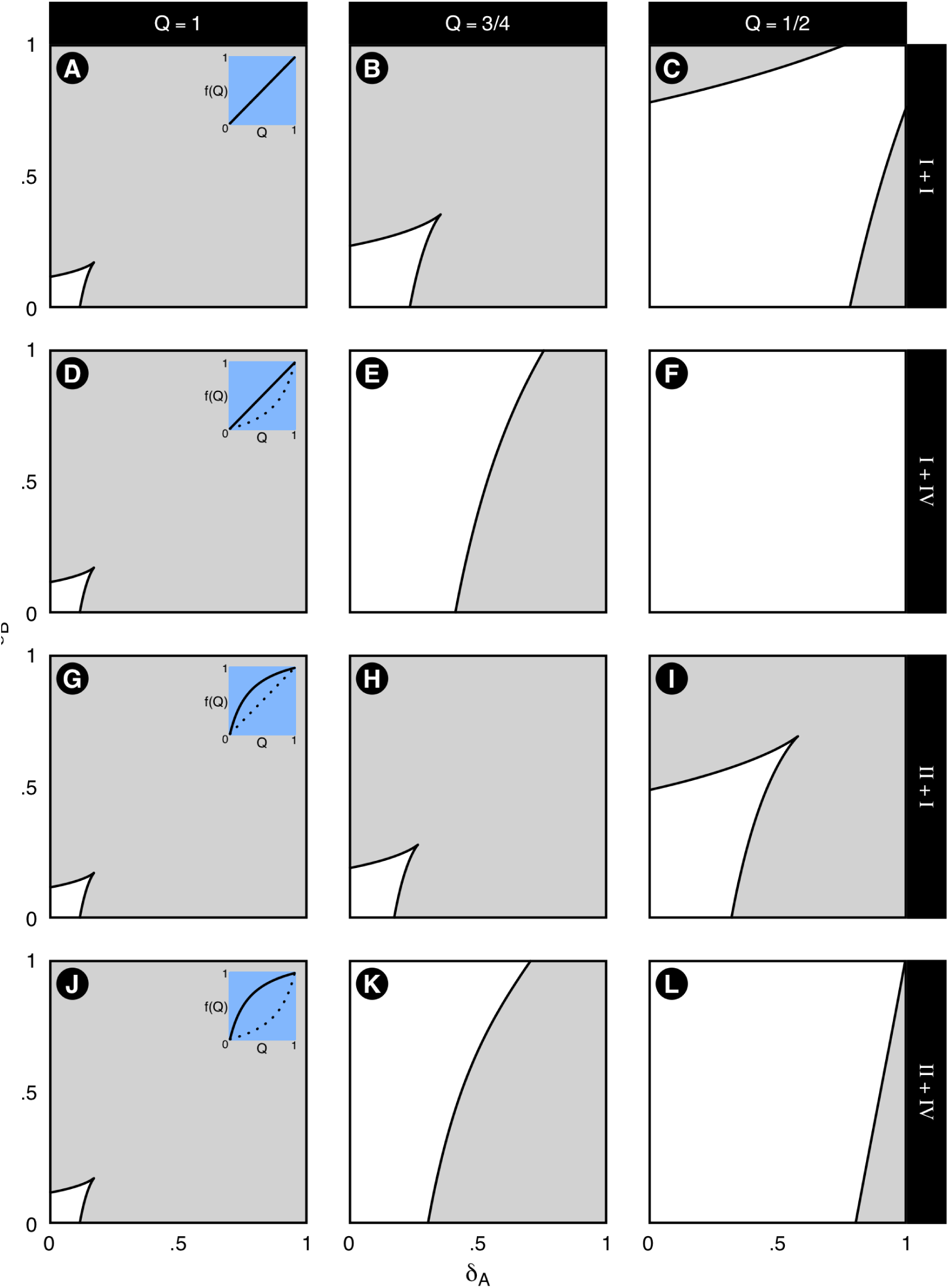
Parameter-space plots showing the boundaries of coexistence for equations 3 and 4 as a function of the strength of reproductive interference suffered by species *A* (δ _*A*_) and *B* (δ _*B*_). Regions where coexistence is possible are denoted in white. Panel rows show different combinations of functional responses for *A* and *B* (*A* is given first, then *B*), which can be linear (type I, *λ*_*i*_ = 1 and *µ*_*i*_ = 1), anthropophilic (type II, 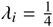 and *µ*_*i*_ = 1) or zoophilic (type IV *λ*_*i*_ = 4 and *µ*_*i*_ = 1). Panel columns show different levels of host availability.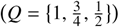 Small interior plots show how the functional responses differ in each of the rows, with the response for *A* shown as a solid line and that for *B* as a dotted line. The first column is the baseline, where *f*_*A*_(*Q*) and *f*_*B*_(*Q*) both equal 1, and the dynamics of Kishi and Nakazawa’s (2013) model (equations 1 and 2) are retrieved. In all cases, the possibility of coexistence increases as host availability is reduced. In panels E, F, H, I, K and L the functional responses are asymmetric and *Q* < 1. Here species *B* is able to suffer higher costs of reproductive interference while still coexisting with *A*. Unvaried parameter values used are given in Table 2, otherwise 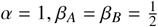.

The interactive effect of reproductive interference and larval competition on the potential for coexistence are summarised in Figure 2. A scenario where species share linear functional responses (white regions) is contrasted with one where species *A* is anthropophilic and *B* is zoophilic (blue regions). Increasing the strength of larval competition decreases the possibility of coexistence, as does increasing the strength of reproductive interference (down the rows). Decreasing the proportion of human hosts available will increase the possibility of coexistence (across columns), as it will reduce the encounter rates between two species. In the asymmetric case (blue regions), as humans become rarer, the zoophilic species is able to suffer a far greater penalty from larval competition and still coexist with an anthropophilic species. This is due to the zoophilic species utilising human hosts less and, therefore, encountering fewer heterospecifics. The discrepancy is greatest in panels H and I, where the cost of reproductive interference is greatest.

**Figure 2:**
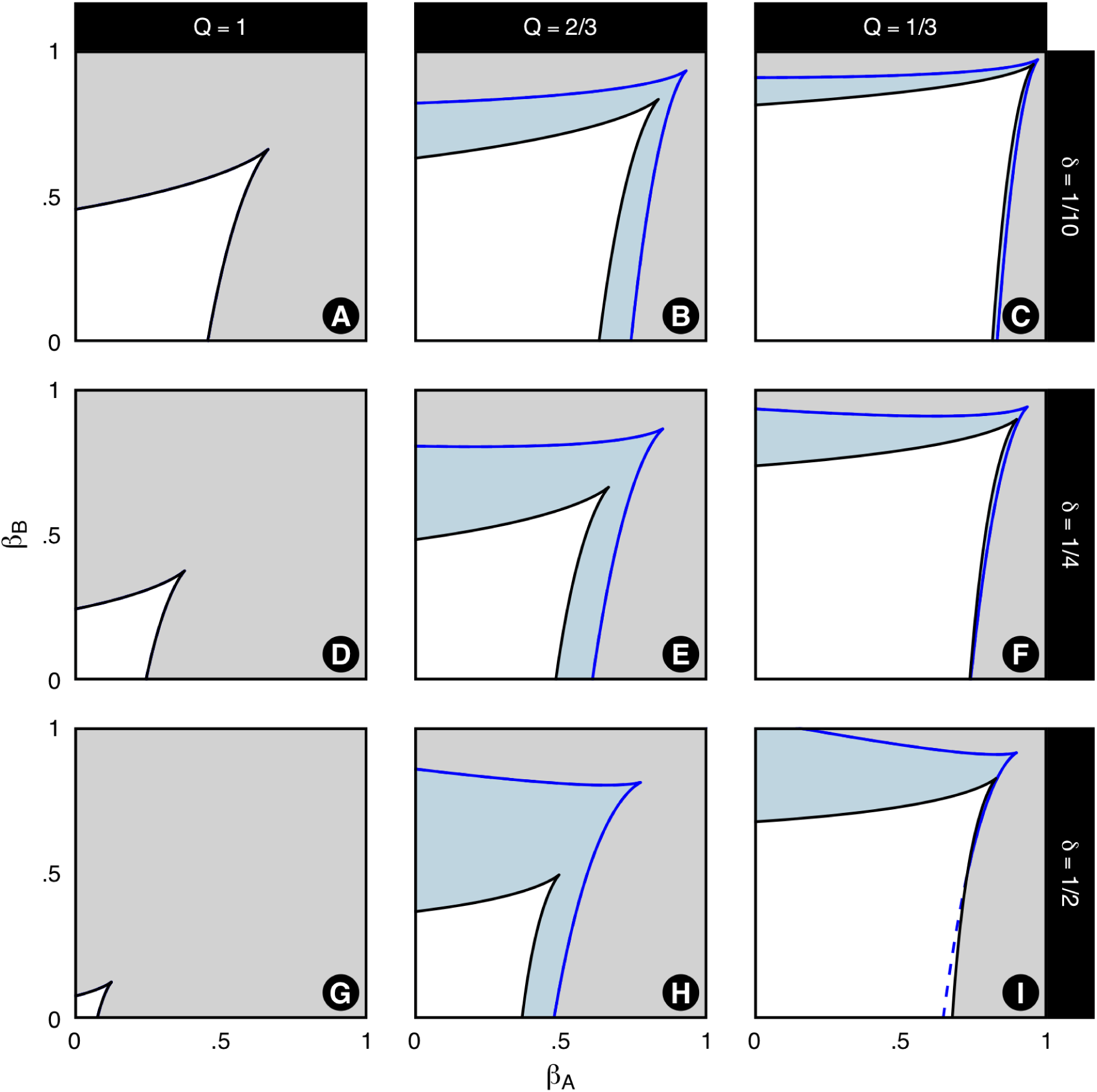
Plots showing how the boundaries of coexistence change as a function of the strength of larval competition suffered by species *A* (*β*_*A*_) and *B* (*β*_*B*_). Panel rows show different levels of reproductive interference for both species 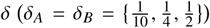 while panel columns show different shared host availability 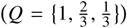 The black line and white regions denote the boundary for a scenario where both species have a linear functional response (both type I, *λ*_*i*_ = 1 and *µ*_*i*_ = 1) and the blue line and shading for a scenario where species *A* is highly anthropophilic (type II, *λ*_*A*_ = 4 and 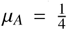) and *B* highly zoophilic (type IV, 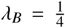 and *µ*_*B*_ = 4). Both cases show that coexistence is possible for greater levels of larval competition when reproductive interference is lower. However, when there are asymmetries in functional responses (blue regions), the species which is disinclined to use the shared host will be able to experience higher levels of larval competition as it suffers a reduced cost from reproductive interference. This is most noticeable in panels H and I, where the cost of reproductive interference is highest. Overall, there is a greater potential for coexistence in the asymmetric functional response cases.

Figure 3 shows the zero-net-growth isoclines derived from equations 5 and 6 for different levels of female sterilisation of species *A* (*ϵ*_*A*_) in the rows and *B* (*ϵ*_*B*_) in the columns. Each panel gives the isoclines for a scenario where larval competition is limited by conspecifics (*α* > *αβ*, solid lines) and heterospecifics (*α* < *αβ*, dotted lines). In panel A, coexistence is not possible as the aggregate costs of reproductive interference and larval competition are greater than the strength of within-species competition. In panels along the diagonal (where *ϵ*_*A*_ = *ϵ*_*B*_), as fewer females of both species contribute eggs, the effects of intraspecific larval competition are diminished to the point where coexistence is possible (panels A, E, I). However, this is only the case when the system is limited by intraspecific larval competition (solid lines). Off-diagonal, coexistence is not possible when discrepancy between *ϵ*_*A*_ and *ϵ*_*B*_ is greatest (panels C and G). However, for the intermediate differences in sterilisation shown in panels F and H, coexistence is possible. Counter-intuitively, in panels F and H the species with *more* sterilised females has a higher predicted coexistence density. This only occurs when larval competition is self-limiting (solid lines), suggesting that this counter-intuitive benefit to the sterilised species is unique to these circumstances.

**Figure 3:**
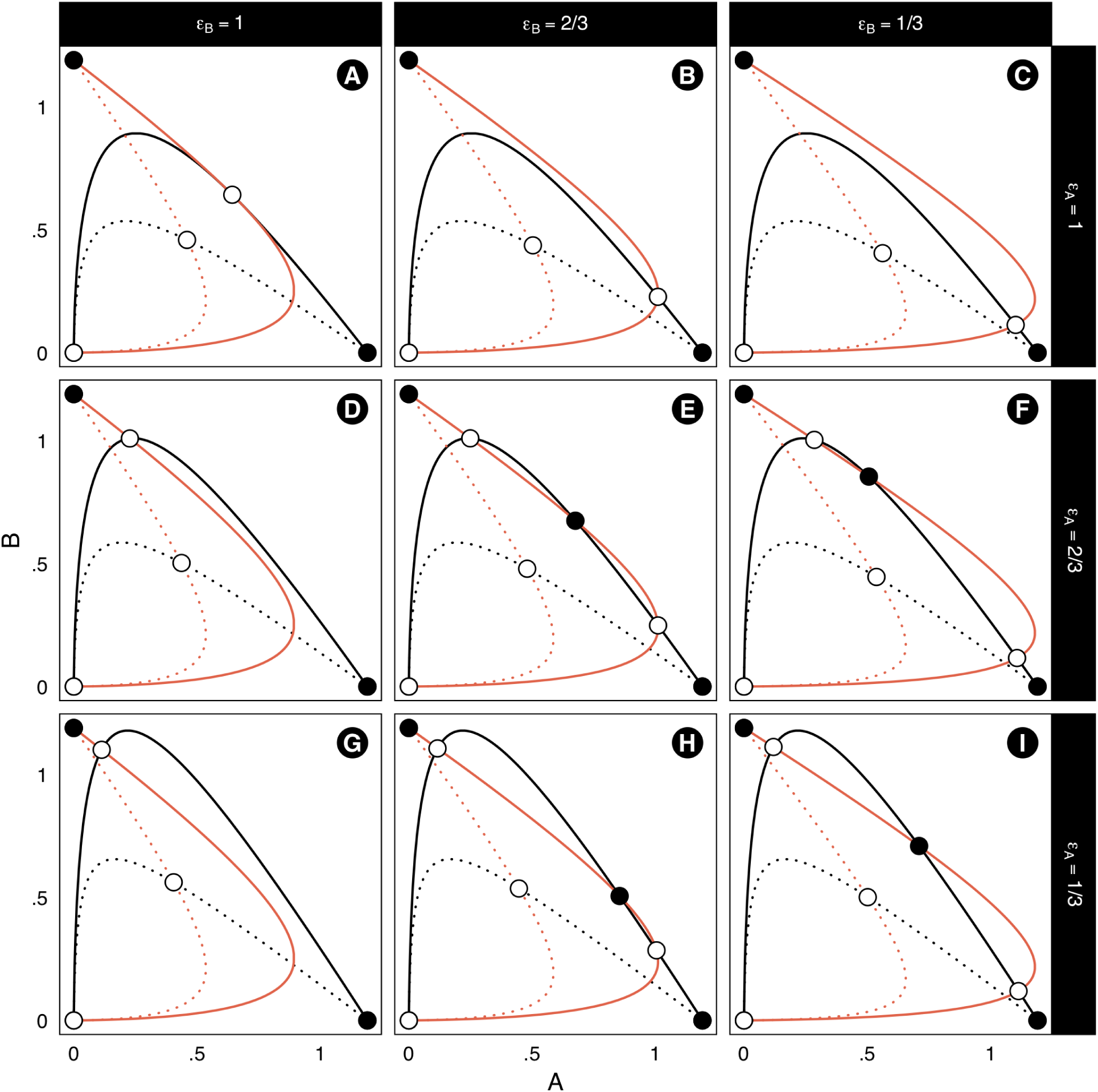
Zero-net-growth isoclines for species *A* (black) and *B* (red) derived from equations 5 and 6. Solid black points denote stable equilibria, and open circles unstable equilibria. Facets correspond to proportions of females of species *A* in the columns 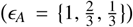 and *B* along the rows 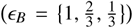 going on to contribute eggs (and therefore larvae) to the process of density-dependence after heterospecific matings. The strength of larval competition is varied within each panel (with *β*_*A*_ = *β*_*B*_). The solid line shows the case where larval competition is self-limiting (*α* > *αβ*) and dotted lines it is limited by heterospecifics (*α* < *αβ*). Along the diagonal the proportion of sterilised females is increased for both species symmetrically, introducing the prospect of coexistence (it is not possible in the initial case with no sterilisation). Off-diagonal, coexistence is possible when intraspecific competition is limiting (solid lines), but the coexistence state is biased toward the species experiencing **more** sterilisation (panels H and F). Unvaried parameter values are given in Table 2, otherwise 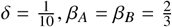 (solid line) or 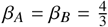 (dotted line).

### 3.2 Epidemiology

In the previous section, we demonstrated that the position and stability of stable-state vector populations is, in part, determined by the process of reproductive interference and host availability (through *δ*_*i*_ and *f*_*i*_(*Q*)). Notably, we have shown that the system could yield either coexistence or single-species dominant states. It is therefore necessary to examine how vector population size affects disease spread so that this can be related to the different configurations of the vector populations. Moreover, the functional responses to hosts will also alter the biting rate on human hosts (Yakob, 2016), suggesting that the alternate-stable states could be epidemiologically non-equivalent. The vector dynamics in this section are described by equations 3 and 4; we do not include examples where vector population dynamics are described by equations 5 and 6. This is because equations 5 and 6 do not include functional responses to hosts, only female sterilisation. In the absence of functional responses, there is no feedback between the process of host selection, reproductive interference and epidemiology.

In Figure 4, the panel backgrounds show how the potential for a disease outbreak (*R*_0_) changes with vector abundance. Overlaid are the zero-net-growth isoclines corresponding to the functional response combinations and host availability for each panel. In the baseline case (the first column), humans are the only hosts, which means it does not matter which functional response the vectors exhibit. In panels A-C, both species have linear functional responses, so *R*_0_ is diminished symmetrically as as human hosts become rarer. In panels E, F, H, I, K and L, the species do not share the same functional responses and humans are not the only hosts (*Q* < 1). This leads to asymmetric values of *R*_0_ depending on which vector is most abundant. This is expected, given that a zoophilic vector will contribute less to human disease than an anthropophilic one. What has not yet been appreciated is that this same process – mediated by reproductive interference - will additionally determine the location of the stable and unstable vector states. For instance, in panels K and L, a stable coexistence point is introduced which falls within the disease outbreak range. A coexistence point would not be introduced if there was no feedback between host-availability and reproductive interference.

**Figure 4:**
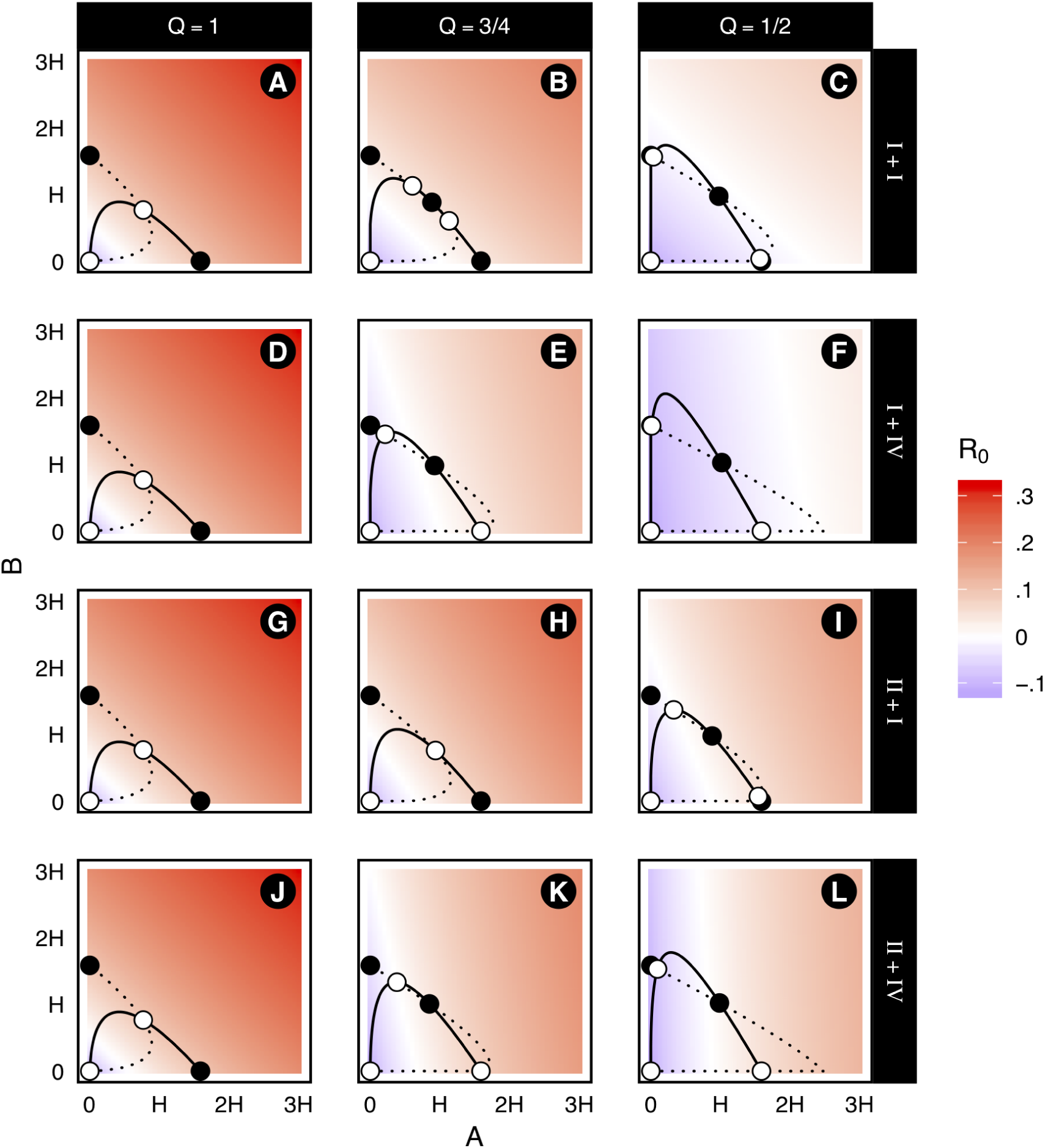
Surfaces exploring the effect of functional responses to hosts on the disease outbreak potential of a two-vector system. The x-axis shows the equilibrum population size of species *A*, and the y-axis species *B*. Axes are given as multiples of the human host population, up to a 3:1 ratio of vectors to hosts. Whether the disease-free equilibrium is stable (*R*_0_ < 0) or unstable (*R*_0_ > 0) across the vector state-space is given by the background colour (blue disease would die out, red it would take-off). Columns denote changes in the availability of hosts 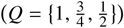 and rows different combinations of linear (type I, *λ*_*i*_ = 1 and *µ*_*i*_ = 1), highly anthropophilic (type II, 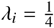 and *µ*_*i*_ = 4) and highly zoophilic (type IV, *λ*_*i*_ = 4 and 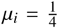) functional responses (first giving the response of *A*, then *B*). Overlaid are the zero-net-growth isoclines (from equations 3 and 4) for the corresponding vector system, with solid lines those of *A* and dotted lines for *B*. Stable-states are denoted with black points, unstable with white. In the first row, where both species respond linearly to host availability, the region over which *R*_0_ > 0 diminishes symmetrically. When there are asymmetries in the functional responses and humans are not the only available hosts (panels E, F, H, I, K and L), *R*_0_ will be lower when the system is biased towards the zoophilic vector and vice-versa. The isoclines (from equations 3 and 4) show the feedback between the functional responses and reproductive interference; changes in host availability can change the location and nature of stable-states, which in turn fall on different values of *R*_0_ (panels K and L). Unvaried parameter values are given Table 2, otherwise 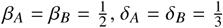 and 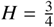

We were motivated to use stochastic simulations of our differential equations to gain a more nuanced understanding of how functional responses to hosts affect the *realised* stability of the vector system, as well as the consequences of multi-stable population dynamics for disease spread. However, exploratory simulations indicated that for all but very small time delays (τ), cyclic or chaotic dynamics would occur. Indeed this was the case when using a time delay informed by empirical observations of *Aedes* development times. For this reason, we assume τ = 0, as highly non-linear dynamics are not the focus of the current study. We suspect that these non-linear dynamics occur because of the linear form of density-dependence in our model; other delay-differential equation models of *Aedes* population dynamics (e.g. Dye (1984)) which use more complex forms of density-dependence do not yield cyclic or chaotic dynamics so readily. They do, however, sacrifice analytic tractability with regards to multi-species modelling.

Figure 5, depicts scenarios with linear/linear (panels A-C) and anthropophilic/zoophilic (panels D-F) combinations of functional responses to hosts. The availability of hosts is changed across columns. The system is simulated to give both the ecological (main plot) and epidemiological (sub-plot) stochastic dynamics. The parametrisation of these scenarios is the same as panels A-C and J-L in Figure 4, where the deterministic isoclines are shown. The local asymptotic stability used to determine the nature of single-and multi-species states identified in Figure 4 is insufficient to describe the full system dynamics. While the states found by the isocline analysis in Figure 4 can be visually identified, there are different probabilities of each state being occupied. The state where the anthropophilic species is dominant is rarely occupied in panels E and F, reflective of the fact that it incurs a greater cost of reproductive interference and is unlikely to out-compete a zoophilic competitor. In the sub plot of panel E, epidemiological dynamics (disease extinction versus disease outbreak) are bimodal, reflecting the transition in the vector state-space. Coexistence with an anthropophilic vector has more severe epidemiological consequences than the single-species dominant state of the zoophilic host, as anthropophilic vectors will seek out rare humans and increase disease transmission.

**Figure 5:**
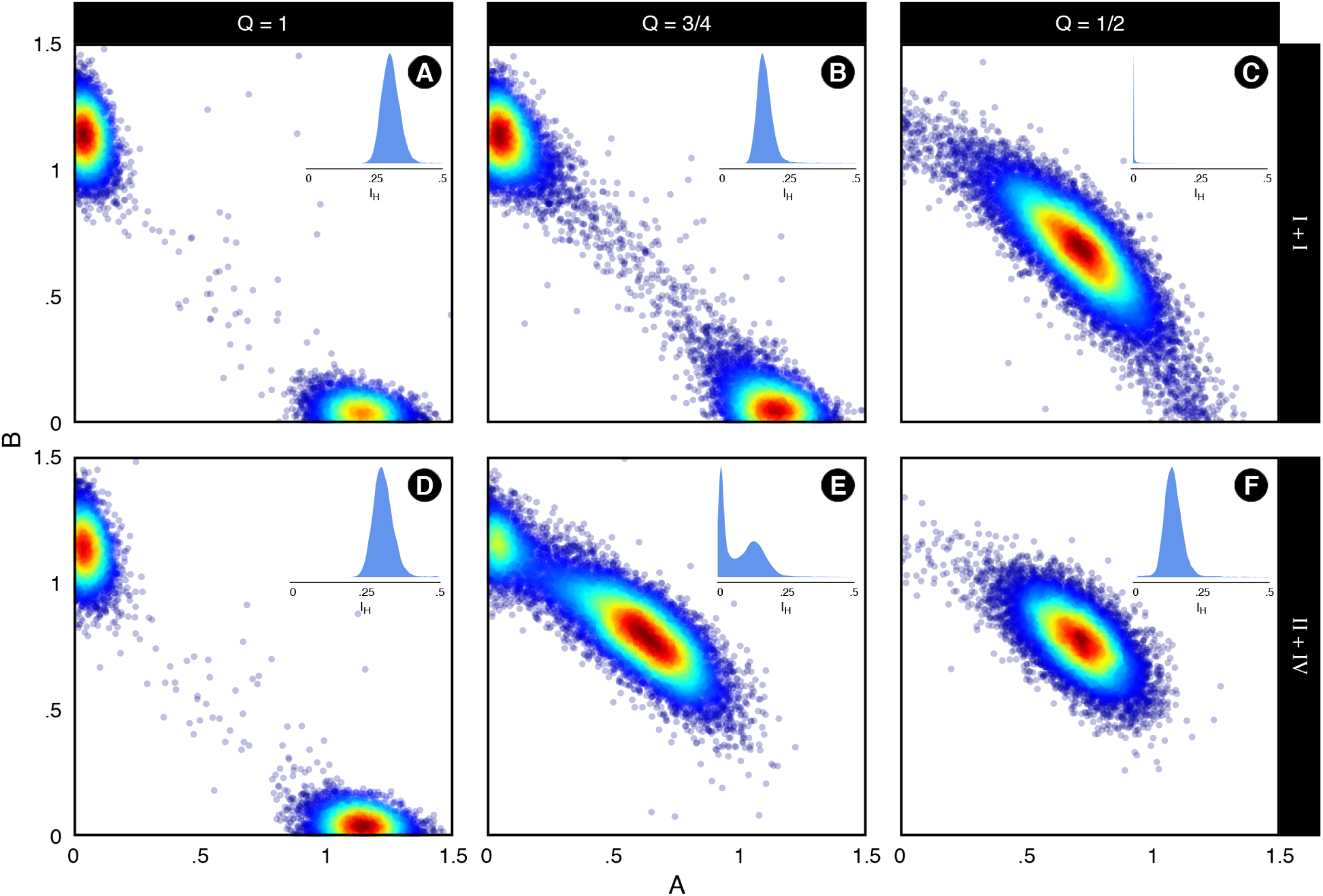
Aggregated results of 100, 1000-iteration stochastic simulations of the multi-vector system with a step-size (*dt*) of 1 × 10^-3^. Scatter-plots of the vector state-space (hot colours indicate higher frequency of occurrence) are shown with density plots of the number of infected humans (*I*_*H*_) shown in the upper right of each facet. We examine are two functional response scenarios. Panels A-C show a “null” case where both have linear (type I, *λ*_*i*_ = 1 and *µ*_*i*_ = 1) responses. Panels D-F show a case where *A* is highly anthropophilic (type II, *λ*_*A*_ = 4 and 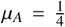) and *B* is highly zoophilic (type IV, 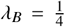 and *µ*_*B*_ = 4). Parameter values are the same as the first and fourth rows in Figure 4, so the deterministic dynamics would be those given by the isoclines in this plot. Observable are the discrepancies between the possible stable states that *could* be occupied and the *realised* states in the stochastic simulations. In panel E, there are frequent state transitions between the *B*-dominant and coexistence states. This corresponds to a diffuse pattern in the number of infected humans, which show a range of values tracking the abundance of the anthropophilic host (sub-plot, panel E). It is notable that the *A*-dominant state is never occupied in this panel. In the final column, the coexistence state becomes the most likely state to be occupied (single-species states are rarely visited), however this corresponds to drastically different epidemiological dynamics. As *A* is anthropophilic in panel F, a coexistence state with this species will allow for disease transmission, whereas it will not for the linear case in panel C. Unvaried parameter values are given Table 2, otherwise 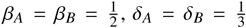 and 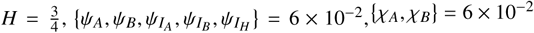 and 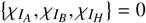.

Figure 6 shows how the distribution of infected hosts (across many stochastic simulations) changes as humans become rarer (i.e *Q* is reduced). Across columns the strength of reproductive interference is increased, while rows denote different combinations of functional responses. In all panels, when only human hosts are available (*Q* = 1), the distribution of infected humans is the same as functional responses cannot alter biting rates. In panels A-C, where both vectors have a linear functional response to hosts, the number of infected hosts declines with host availability, doing so more steeply when reproductive inference is stronger (panels B and C). This outcome is intuitive; as hosts become rarer, the vector will bite human hosts less and there will be fewer infections. When functional responses are not the same (panels D-L), there is a non-linear relationship between host availability and the distribution of infections. The combination of reproductive interference and host selection lead to a concomitant relationship between ecology and epidemiology. As humans become rarer, the costs from reproductive interference are reduced and the biting rate on hosts lessened. This reduced competitive cost can facilitate the coexistence of vectors in circumstances where the zoophilic vector would have otherwise dominated, leading to a higher *R*_0_ than the linear/linear case. This non-linearity is most clear in panels H and L. In panels H, I and L, bimodal distributions of infected humans are observed as the system transitions between single-species dominant and coexistence states (the same dynamics are observed in Figure 5, panel E). The state where the more anthropophilic species is dominant is rarely occupied as it suffers a greater cost from reproductive interference, suggesting that it is very unlikely that the zoophilic species will be outcompeted.

**Figure 6:**
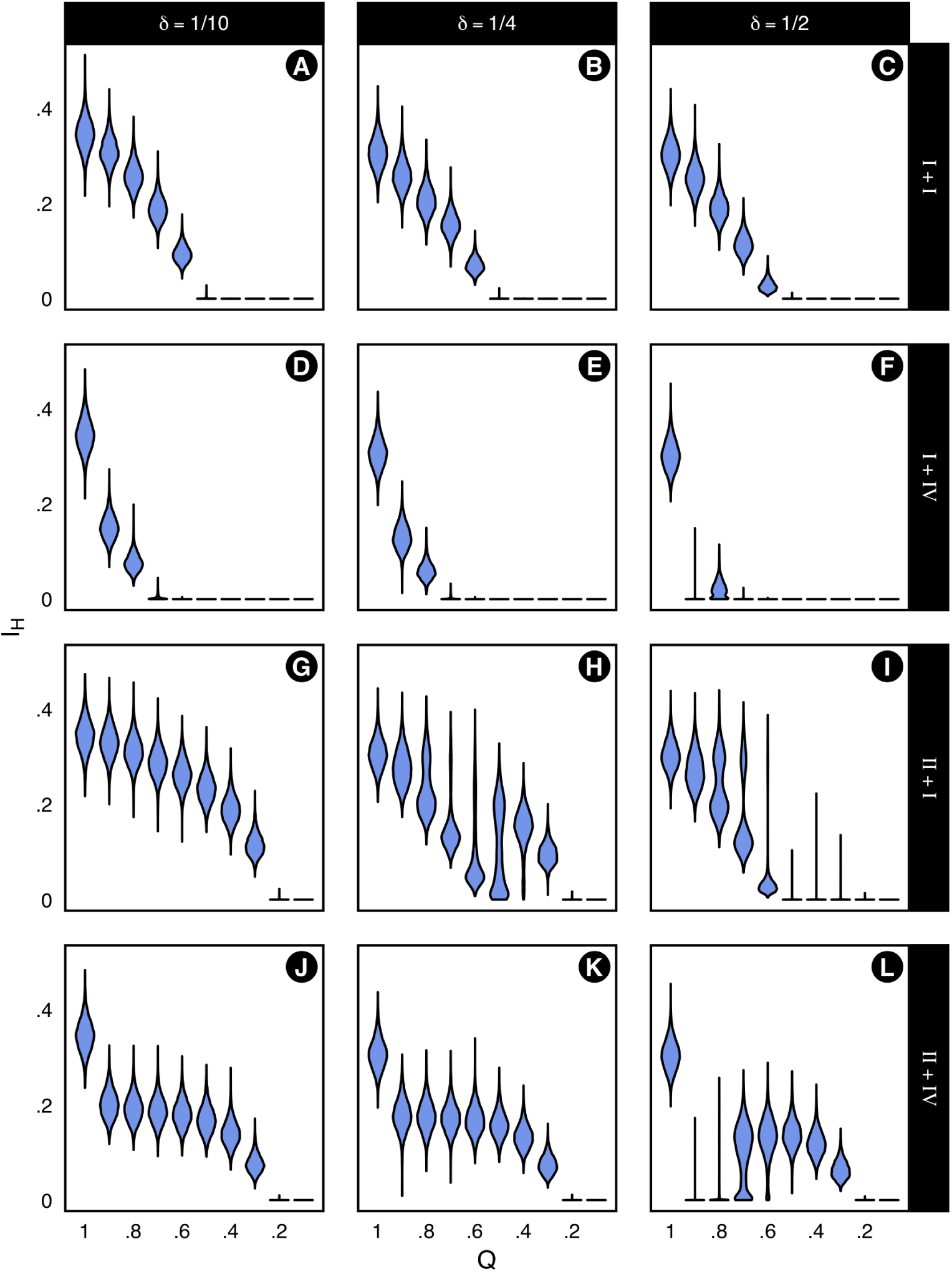
Violin plots demonstrating how changes in host availability (*Q*, from 1 to 0.1 in increments of 0.1) affect the distribution of infected humans (*I*_*H*_). Each violin is a summary of 100, 1000-iteration stochastic simulations of the multi-vector system with a step-size (*dt*) of 1 × 10^-3^. Functional responses are varied in the rows, with scenarios a combination of linear (type I, *λ*_*i*_ = 1 and *µ*_*i*_ = 1), highly anthropophilic (type II, *λ*_*A*_ = 4 and 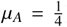) and highly zoophilic (type IV, 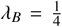 and *µ*_*B*_ = 4) functional responses. The response of species *A* is given first, then *B*. Columns show different strengths of reproductive interference, where the costs are the same for both species 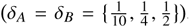 In the first row, the number of infected humans decreases with host availability (both hosts have linear responses). In the asymmetric cases, dynamics can be highly non-linear. Panels H and L show an initial reduction in the number of infected humans before infections counter-intuitively begin to increase with decreasing host availability. In panels H, I, and L, human infections are bimodally distributed for some values of *Q*. This is indicative of shifts between single-species dominant and coexistence states. The epidemiology of these dynamics are contingent on which species is dominant (coexistence with an anthropophilic will result in a greater transmission potential than a zoophilic dominant state). Effects are most severe when the cost of reproductive interference is larger (i.e. across rows). Unvaried parameter values are given Table 2, otherwise 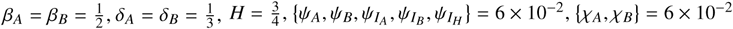 and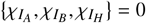.

## 4. Discussion

We present an *Aedes*-bespoke theoretical framework for examining the roles of two mechanisms of competition in mosquito population dynamics and epidemiology. In our analysis, we have explored two candidate mechanisms that could explain the inconsistent patterns of co-occurrence and competitive exclusion observed in wild populations of *Ae. aegypti* and *Ae. albopictus*: larval competition and reproductive interference. These mechanisms were embellished with *Aedes*-bespoke features relating to these forms of competition: functional-response mediated encounter rates and sterilisation of females by copulation with heterospecific males. We examined the consequences of these ecological nuances on the epidemiology of a theoretical vector-borne disease.

### 4.1. Ecological insights

#### 4.1.1. Functional responses to host availability

Our findings suggest that population coexistence and competitive exclusion could be driven by differences in the availability of a shared host if competitors suffer from reproductive interference and have divergent functional responses to hosts. This is because *Aedes* use hosts to find mates (Nelson, 1986; Yuval, 2006), suggesting that *Ae. aegypti* and *Ae. albopictus* will only encounter heterospecifics (and incur the cost of reproductive interference) at shared hosts. We show that when the functional responses of both species are linear the effects of reproductive interference are symmetrically scaled and the possibility of coexistence is increased-even when the associated costs are high. This is due to the encounter rate with heterospecifics being reduced in tandem with humans becoming rarer. More interesting are cases where the vectors have *different* functional responses. In these cases, as host availability is reduced, the possibility of coexistence varies asymmetrically with the costs associated with reproductive interference; the species with the more zoophilic functional response can suffer higher costs of reproductive interference while still coexisting with the anthropophilic species. This is intuitive, as a smaller proportion of the zoophilic population will be exposed to heterospecifics and incur the associated costs.

Also notable is the indirect effect of reproductive interference on larval competition. While the functional responses do not *directly* affect the process of reproductive interference, it will still influence the costs associated with larval competition. With linear functional responses, reducing human host availability increased the strength of larval competition (*β*_*i*_) that can be suffered while still permitting the species to coexist. However, we show that when functional responses are asymmetric, zoophilic species can suffer higher levels of larval competition while still coexisting with the anthropophilic species. Overall, we find that higher costs of reproductive interference reduce the possibility of coexistence for high coefficients of larval competition.

Kishi and Nakazawa (2013) provided a rigorous analysis of Kuno’s (1992) model, and gave valuable insights into the processes of reproductive interference and how it might interact with Lotka-Volterra competition. When functional responses are linear, our results corroborate this finding as the function describing reproductive interference will linearly scale with the exposure to heterospecific mosquitoes. Our work has expanded this to investigate non-linear functional responses, demonstrating that functional responses can drive significant asymmetries in the levels of competition experienced by each species, leading to different patterns of coexistence and exclusion, even when the strength of reproductive interference and larval competition are the *same*. This is contingent on the shared host not being the only available host. For instance, if a lab- or field-study determined the same coefficients of larval competition and reproductive interference for both *Ae. aegypti* and *Ae. albopictus*, then our model predicts that, in practice, the availability of the hosts could result in these costs manifesting in different competitive outcomes if the species have different functional responses. These discrepancies between the measured costs of competition and the *realised* costs under different host regimes is crucially important for those studying, analysing (Juliano, 2010) and reviewing (Juliano, 2009) the competitive effects found in *Aedes* lab, semi-field and field studies. In order to correctly interpret study findings (e.g. which competitor is superior) the environmental context must also be accounted for. This is in addition to already understood dependencies, such as resource (Reiskind et al., 2012), population (Leisnham and Juliano, 2009) and habitat (Rey et al., 2006) mediated differences in competitive outcome. Indeed further complications are likely to include strain-specific responses to host availability (it is unlikely that all *Ae. aegypti* and *Ae. albopictus* populations will share the same host preferences).

#### 4.1.2. Sterilised females

Our model also explored the effect of sterilisation of female mosquitoes by matings with heterospecific males (Tripet et al., 2011) on patterns of coexistence and exclusion. This is an another indirect effect of reproductive interference on larval competition; when both species become sterilised, we find that coexistence is promoted as larval density-dependent competition is alleviated. The findings for asymmetric cases (where levels of sterilisation in the two species differ) depend on whether the system is limited by interspecific or intraspecific larval competition. Counter-intuitively, we find that the sterilisation of females by heterospecific matings lead to *higher* equilibrium densities of the species experiencing higher levels of sterilisation when coexistence was possible. However, this occurs only when the population is limited by intraspecific larval competition. A comparable phenomena is observed with the superior efficacy of late-acting lethal control strategies (e.g. GM mosquitoes) compared with early-acting sterilising strategies (e.g. sterile insect technique) (Phuc et al., 2007). Theory suggests that removing the density-dependent competition by “skimming-off” excess individuals will increase the reproductive rate of the species. In the *Aedes* case, alleviating the larval competition in the shared pools would increase survivorship and lead to an increased reproductive rate for the sterilised population (relative to the reproductive interference free situation). Similar models, such as those for tsetse fly control (Rogers and Randolph, 1984) suggest a similar increase in the equilibrium populations sizes would be observed under sterile insect releases. The unifying assumption is that populations are density-limited.

The counter-intuitive increase in equilibrium population density arising when mortality (or in our case sterility) is termed a “hydra-effect”. Abrams (2009) outlined three candidate mechanisms which could cause this, with one particularly relevant for our model; “the temporal separation of mortality and density-dependence”. While the process of sterilisation in our second model (equations 5 and 6) does not describe an increase in mortality *per se*, the outcome is the same in that larvae experience reduced density-dependent regulation. Abrams (2009) also identify that in order for this mechanism to produce a hydra-effect, then there must be over-compensatory density-dependence which is absent from our model as our within-species competition is only compensatory. McIntire and Juliano (2018) provided empirical support for Abrams’s (2009) temporally-separated mortality hypothesis by rearing cohorts of *Ae. albopictus* in the lab and exposing them to a regime of either early and late-acting mortality. They found that early-acting mortality resulted in higher adult densities. This suggests that the predictions of models in the vector-control literature are correct in cautioning against sterile insect techniques when species are limited by strong over-compensatory density-dependent processes, and gives real-world context to the predictions of our model.

The sensitivity of these conclusions to the system being limited by within or between-species competition is clear; if conspecifics exert a greater cost in terms of within-species larval survivorship, then fewer conspecifics will allow them to do better. However, if the sterilised species suffers a greater cost from heterospecific larvae than conspecific larvae, this process will not confer the same benefit.

### 4.2. Epidemiological consequences

We have extended the epidemiological insights provided by Yakob’s (2016) research into functional responses to host availability in two ways. First, the same framework can be applied to multi-vector systems with different host preferences. Second, host selection has *ecological* consequences for *Aedes*. Indeed, we have shown the ecological and epidemiological processes to be inextricably linked with regards to reproductive interference. These effects can be both detrimental and beneficial for vector-borne disease transmission. Our exploration of the disease potential of the multi-vector state-space showed that the potential for outbreaks varies drastically with different population sizes of zoophilic and anthropophilic vectors. The process of reproductive interference determines the region of this state-space likely to be occupied by vector populations, leading to a complex feedback of the two processes; an anthropophilic vector is more competent at transmitting disease, but will be more likely to be excluded by reproductive interference. The reverse may also be true however, as reducing human host availability may, through diminished reproductive interference, balance other costs (such as asymmetric effects in larval competition) and promote coexistence between zoophilic and anthropophilic species.

Our stochastic simulations tie-in with our deterministic findings by describing the *realised* stability of the system, not solely the asymptotic stability precisely at the equilibrium. Evident from our simulations is that the deterministic predictions do not fully demonstrate dynamical differences between the scenarios. In our example, the multi-stable states are not equivalently likely, with some states more likely to precipitate a state-change than others. Stochastic processes will more easily shift the system between states of comparable stability, and may prohibit the occupation of some states as they are simply too unlikely. Our simulations also show that changes in host availability and functional responses can drastically alter system stability, and promote the occupation of certain states. In the example, we show that asymmetries in functional responses can result in drastically different dynamics, which correspond to very different epidemiological outcomes; as states transition from single-species dominant to multi-vector states the system can fall in a region of the state-space where an outbreak is possible. In short, selecting which host to bite determines the level of reproductive interference suffered, as well as the outbreak potential in humans. This leads to a highly non-linear relationship between host availability and the outbreak of infection when the species have different functional responses.

### 4.3. Insights, assumptions and extensions

Bonsall et al. (2010) analysed a model of *Aedes* larval competition without the process of reproductive interference, instead including the effect of sterile or GM (late-acting lethal) mosquito releases targeted at one of the species. They derived non-linear isoclines, driven by one species experiencing population control. Our work raises an important question with regards to population control; if both species experience reproductive interference, then releases targeted at one species will almost certainly affect the non-target population. We advocate the exploration of the interaction between reproductive interference and targeted-population suppression techniques, as the effects on predicted coexistence and exclusion could be profound.

In our model we use a frequency-dependent function for describing the process of reproductive interference; increasing the relative frequency of heterospecifics to conspecifics scales the reproductive rate. Kyogoku and Sota (2017) present a model of reproductive interference where heterospecific encounters are governed by a density-dependent function (rather than the frequency dependent function used here and by Kuno (1992)). Their model is noteworthy in that it can produce a more diverse range of patterns of coexistence and exclusion (for instance, unlike our frequency-dependent function, invasions from rare are possible if the *density* of heterospecifics is sufficiently low). We elected not to use this model as a our frequency-dependent function is recovered when the population densities are very high. However, the model presented by Kyogoku and Sota (2017) provides interesting insights in several cases. For instance, if there are very few vectors and very large numbers of hosts, then encounter rates may be diluted and the effects of reproductive interference drastically diminished. We recommend this as a focus for further exploration in relation to the phenomena we describe here.

The exact form of larval density-dependant competition in *Aedes* is still debated and difficult to measure in wild populations (Legros et al., 2009). Here we have assumed that a linear function is an adequate description of the process (i.e. the logistic-assumption holds, with each larvae contributing equally to density-dependence). Examining the interaction between non-linear forms of density-dependence and the competitive processes outlined in this research is a logical and worthwhile extension. Given the links between over-compensatory density-dependence and the Hydra effect (Abrams, 2009; McIntire and Juliano, 2018), exploring different forms of population feedback will be valuable in elucidating the role of population regulation in increasing equilibrium densities of sterilised species with competitive interactions.

## 5. Conclusion

Kuno’s (1992) model of reproductive interference and Yakob’s (2016) study of vector functional responses gave independent insights into two crucial processes. The models presented in this paper are unique in uniting these insights into a model of *Aedes* competition. We have elucidated the relative roles of reproductive interference and larval competition, showing that host selection can allow these processes to be “traded-off”, allowing for coexistence where it would be difficult to predict from practical studies. We have also shown that these ecological and epidemiological processes are inextricably linked, with host selection determining the costs of reproductive interference (potentially leading to vector coexistence) as well as biting rate. In doing so, we have added to the complex range of candidate mechanisms which could explain variable patterns of exclusion and coexistence between *Aedes* mosquitoes.

## 6. Acknowledgements and author contributions

RSP was supported by a NERC studentship (NE/L002612/1) and is a CASE student with the Pirbright Institute. MBB is supported by two BBSRC grants (BB/H01814X/1 and BB/L00948X/1). RSP developed the concept and ideas, with advice and input from MBB. RSP carried out the analysis and wrote the manuscript. MBB reviewed the manuscript and contributed to the final document. The authors would like to thank Tom Brewer for reviewing the document thoroughly and making detailed recommendations and suggestions.

## Appendix.1. Isocline derivation equations 3 and 4

Solving equations 3 and 4 for 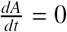 and <u>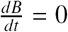</u> respectively and factorising yields two expressions

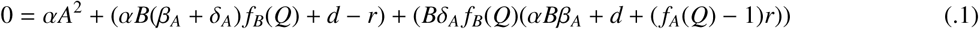

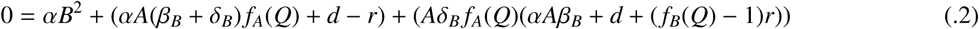

The solutions to equations .1 and .2 are simply the positive solution of the quadratic formula. Substituting the coefficients for equations .1 and .2 the isoclines for the state of species *A* and species *B* are obtained

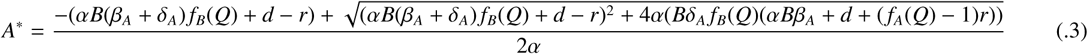

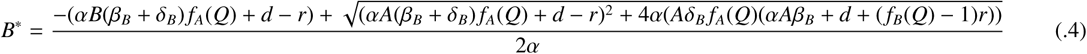

The necessary conditions for coexistence can be examined by substituting equation .3 into .4 and vice-versa yields. This yields a cubic polynomial for each species, and is included in the supplementary Mathematica file. The discriminant of this expression reveals the nature of the roots of the polynomial.

## Appendix .2. Derivation of R_0_

The *R*_0_ of this multi-species system can be calculated from the following. The Jacobian matrix of the linearised system is given below, based on equations 7 to 9

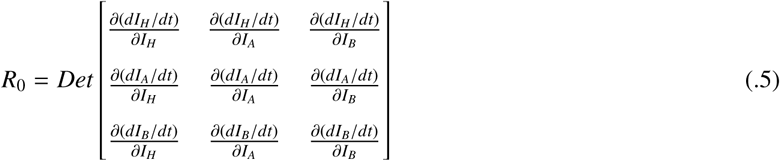

Then, calculating the partial derivatives

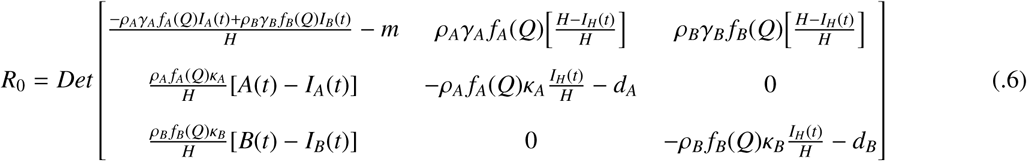

Assuming that the pathogen invades an entirely healthy population (*I*_*A*_(*t*), *I*_*B*_(*t*) and *I*_*H*_(*t*) → 0) and vector population sizes are at equilibrium (*A*(*t*) = *A*^*^ and *B*(*t*) = *B*^*^) then *R*_0_ is

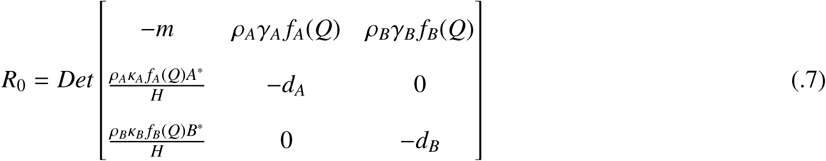

